# An *in vitro* and computational validation of a novel loss-of-functional mutation in PAX9 associated with non-syndromic tooth agenesis

**DOI:** 10.1101/2021.12.31.471205

**Authors:** Tanmoy Sarkar, Prashant Ranjan, Smitha Kanathur, Ankush Gupta, Parimal Das

**Affiliations:** Centre for Genetic Disorders, Institute of Science, Banaras Hindu University, Varanasi; Department of Periodontology, Government Dental College and Research Institute, Bangalore, India; Department of Biochemistry, Institute of Science, Banaras Hindu University, Varanasi, Uttar Pradesh, India; Mouse Cancer Genetics Program, Center for Cancer Research, NCI-NIH, Fort Detrick, Frederick, MD, USA

**Keywords:** Congenital tooth agenesis, PAX9, Novel mutation, Functional analysis, RNA-sequencing

## Abstract

Congenital tooth agenesis (CTA) is one of the most common craniofacial anomalies. Its frequency varies among different population depending upon the genetic heterogeneity. CTA could be of familial or sporadic and syndromic or non-syndromic. Five major genes are found to be associated with non-syndromic CTA namely, *PAX9, MSX1, EDA1, AXIN2* and *WNT10A*. In this study, an India family with CTA was investigated and a novel c.336C>G variation was identified in the exon 3 of *PAX9*, leading to substitution of evolutionary conserved Cys with Trp at 112 amino acid position located at the functionally significant DNA binding paired domain region. Functional analysis revealed that p.Cys112Trp mutation did not prevent the nuclear localization although mutant protein had higher cytoplasmic retention. EMSA using e5 probe revealed that mutant protein was unable to bind with the paired-domain binding site. Subsequently, GST pull-down assay revealed lower binding activity of the mutant protein with its known interactor MSX1. Further RNA-sequencing of PAX9 over-expressed HEK293, identified two potential novel targets, *WNT4* and *WNT7b* those are up-regulated by wild-type PAX9 but not by mutant. These *in vitro* results were consistent with the computational results. The *in vitro* and computational observations altogether suggest that c.336C>G (p.Cys112Trp) variation leads to loss-of-function of PAX9 leading to CTA in this family.

## 1 Introduction

Failure of tooth eruption in oral cavity due to immature termination of tooth development is called congenital tooth agenesis which causes reduction of number of teeth from the normal [for human 32 permanent teeth and 20 deciduous teeth]. Considering third molars around 20% of world population show tooth agenesis although it has been considered as an evolutionary consequence of human jaw size reduction. Excluding third molars the frequency of tooth agenesis varies from 2.6% to 11.3% with significantly higher in males than that of females (Polder et al., 2004; De Coster et al., 2009). Depending upon the severity, congenital tooth agenesis (OMIMID #106600, #604625, #150400, #616724, #617073, #617275 and #313500) in human can be classified into three different forms: Hypodontia (Congenital absence of one to five teeth), oligodontia (Congenital absence of six and teeth), Anodontia (Congenital absence of all primary and/or permanent teeth). Except for third molar, mandibular second premolars show most frequent congenital tooth agenesis (2.91-3.22%), followed by maxillary lateral incisors (1.55-1.78%), maxillary second premolars (1.39-1.61%) and maxillary central incisors are least affected (0.00-0.01%) (Polder et al., 2004; De Coster et al., 2009).

There are five genes that are frequently found to be associated with congenital tooth agenesis, namely Paired box 9 (*PAX9*) (Stockton et al., 2000), Muscle segment homology homeobox 1 (*MSX1*) (Vastardis et al., 1996), Ectodysplasin A (Tao et al., 2006), Axis Inhibition Protein 2 (*AXIN2*) (Lammi et al., 2004), Wingless type MMTV integration site family member 10A (*WNT10A*) (Lammi et al., 2004; Kantaputra and Sripathomsawat, 2011; van den Boogaard et al., 2012). Subsequent studies show that *PAX9* is the major candidate gene associated with familial non-syndromic CTA (Bonczek et al., 2017). Recently *EDAR* and *WNT10B* genes are also found to be associated with non-syndromic tooth agenesis (Arte et al., 2013; Yu et al., 2016). With the availability of high throughput DNA sequencing technique new candidate genes such as *SMOC2* (Alfawaz et al., 2013), *LTBP3* (Dugan et al., 2015), *RUNX2* (Molin et al., 2015), *LRP6* (Massink et al., 2015) and *GREMLIN2* (Kantaputra et al., 2015), have been identified to be associated with non-syndromic or syndromic tooth agenesis.

During the process of tooth development, *Pax9* and *Msx1* transcription factors play an essential role in the establishment of odontogenic potential from dental epithelium to dental mesenchyme. Both the genes expressed in dental mesenchyme and interact with each other to activate one of their known downstream targets *Bmp4*.Which subsequently establishes enamel knot, a major signalling centre that controls further tooth development. Several studies indicated that mutations in *PAX9* disrupt PAX9-MSX1 interaction and also sequence specific DNA binding activity of PAX9.

In this present study, we have identified a novel pathogenic mutation in *PAX9* gene responsible for autosomal dominant type of non-syndromic oligodontia in an Indian family. Pathogenicity of the identified mutation was further validated using functional analysis of biological activity of mutant protein.

## 2 Materials and methods

### 2.1 Recruitment of patient and family samples

All the patients and their unaffected family members belong to a multi generation family of Bengaluru, India was enrolled for the study. Congenital tooth agenesis in individuals was confirmed by orthopantomogram (OPG) followed by clinical examination for ectodermal organ namely hair, nails, skin and teeth. Information regarding sweating, heat tolerance and any other congenital anomalies were collected by interview at Department of Periodontology, Government Dental College and Research Institute, Bengaluru, India. After obtaining written informed consent from all participants, OPGs were collected from seven individuals and 5 ml of peripheral venous blood samples from four affected and one unaffected individual were collected in EDTA vial. Subject consent and study protocol was approved by the Institutional Review Board of Banaras Hindu University (No.Dean/2008-09/484)

### 2.2 Mutation detection and in silico analysis

Genomic DNA was extracted from peripheral venous blood samples. All the coding exons and exon-intron boundaries of *PAX9* and *MSX1* and putative promoter region of *PAX9* were amplified using polymerase chain reaction (PCR) from four affected, one unaffected and one assumed to be affected individuals. DNA extraction protocol, PCR condition and primers were used as described previously (Sarkar et al., 2014). PCR products were treated with exonuclease I and recombinant Shrimp alkaline phosphatase (rSAP) (USB Affimetrix, USA) followed by labeling with ABI Big Dye Terminator V3.1 cycle sequencing kit for Sanger sequencing. Capillary electrophoresis and automated base calling was performed in ABI 3130 Genetic Analyzer and Sequencing analysis software V5.2 (Applied Biosystems, USA) according to manufacturer’s protocol. All the sequences were aligned with available National Center for Biotechnology Information (NCBI) Gen Bank DNA sequence data base using NCBI-Basic Local Alignment Search tool (BLAST). 200 unrelated control chromosomes from 100 individuals (18 females and 82 males) of same ethnic background and within 25 to 30 years of age were examined for this potentially novel variation using automated DNA sequencing.

### 2.3 In silico analysis of identified variation

#### Secondary structure prediction, Homology modeling and Energy minimization

Identified variation was tested for their pathogenic potential using available online software tool namely PolyPhen-2 and MutationTaster (Adzhubei et al., 2010; Schwarz et al., 2014).

Crystal structures of MSX1 protein (PDBID-1IG7) was retrieved from PDB RCSB (https://www.rcsb.org/). All water molecules and hetero-atoms were removed by using Discovery studio visualization software (BIOVIA 2020). (http://accelrys.com/products/collaborative-science/biovia-discovery-studio/visualization-download.php). Secondary structure of wild type and mutant PAX9 was predicted using Psipred tool (Liam et al, 2000) using Pax5 (PBB ID: 1k78.2.C) followed by Homology modeling using SWISS-MODEL followed by Energy minimization using Mod-refiner. Evaluation of the modeled structure was done by PDB-Sum (Laskowski et al, 2018) (http://www.ebi.ac.uk/thornton-srv/databases/cgibin/pdbsum/GetPage.pl?pdbcode=index.html).

#### Protein-Protein and DNA-Protein interaction using Docking

Interaction between PAX9 (wild type and mutant) and MSX1 was carried out PatchDock docking server (Ranjan *et al*., 2021; Ranjan *et al*., 2020). Followed by binding energy and other biochemical parameter calculation using Prodigy web server (Xue et al, 2016) (https://bianca.science.uu.nl/prodigy/) and visualization by PDB-Sum web server via DMPLOT script algorithm.

For DNA-Protein interaction, DNA modeling of e5 probe (CACGGGACAGCCTGAGCGGAACGGTGCTAATCGTGCGGT) was done by DNA sequence done by structure-SCFBio tool (http://www.scfbio-iitd.res.in/software/drugdesign/bdna.jsp). Docking of modeled PAX9 and DNA structure were performed using PatchDock server with cluster RMSD set to 4.0 as the by default. The geometric shape complementarity score (GSC score) calculation and visualized by Chimera 1.5.1 and Discovery studio.

#### Molecular Dynamics Simulation (MDS)

The equilibrium and the molecular dynamic behavior of wild and mutant variants of PAX9 protein was studied by using GROMACS (Hess, *et al*., 2008). GROMACS96 54a7 force field (Ranjan *et al*, 2021) was used for MD simulation study. We added dissolvable water around protein to work with from spc216.gro as a non-elite equilibrated 3-point dissolvable water model in a dodecahedron. At this juncture, we kept the protein in the middle in at least 1.0 nm from the case edges. We utilized steepest descent algorithm for energy minimization in the further steps. Then, we equilibrate the framework by means of NVT ensemble (constant Number of particles, Volume and Temperature) and NPT outfit (constant Number of particles, Pressure and Temperature). Subsequent to accomplishing equilibrium measure, we moved for MD run to 10 ns analysis done by Gromacs tools (i.e. gmx rms, rmsf, gyrate hbond, and sasa respectively) for RMSD (Root Mean Square Deviation), RMSF (Root Mean Square Fluctuation), gyrate for radius of gyration (Rg), H-bond (for intra-protein H-bonds and for H-bonds between protein and water), SASA (solvent accessible surface). Further we used XMGRACE software for data visualization.

### 2.4 Construction of mammalian expression plasmid and site directed mutagenesis

A 1.6-kb full length human PAX9 cDNA clone (GeneBank™, accession number NM_006194.1) containing 5’ and 3’ UTRs was purchased from OriGene Technologies. Using this as a template 1023bp coding region of PAX9 was amplified using forward (5’TAATATGGATCCGAGGAGCAATGGAGCCAGCCTTC3’) and reverse (5’TAATATGTCGACCAGCGCGGAAGCCGTGACAGAATG3’) primers containing BamHI and SalI site respectively. PCR reaction was carried out using Platinum Taq DNA Polymerase High Fidelity (Invitrogen, USA) in ABI Veriti 96 well thermal cycler (Applied Biosystems, USA) programmed with initial denaturation at 94°C for 2 min., followed by 30 cycles of 94°C for 25 sec., 55°C for 25sec., 68°C for 1min. and one final extension at 68°C for 5min. Resulting 1055 bp PCR product was gel-purified using Qiagen Gel Extraction kit (Qiagen Inc., USA) according to manufacturer’s protocol. Purified product was digested with BamHI and SalI restriction endonucleases (TAKARA, Japan) and sub cloned in BamHI and SalI site of pCDNA3.1/myc-His(A) mammalian expression vector (Invitrogen, USA) to generate pCDNA3.1-PAX9-myc-His(A) encoding recombinant PAX9 protein with myc and His tag at C-termini. To create FLAG-*MSX1*-pCDNA3.1V5-His plasmid, CDS was PCR amplified from human full length cDNA clone (Open biosystems, USA) with FLAG sequence flanking at 5’ end of the primer and cloned in Hind III and Xba I site of pCDNA3.1V5-His (Invitrogen, USA). Clones were sequenced with T7 and BGH primer using ABI BigDyeTerminator V3.1 cycle sequencing kit followed by automated sequencing in ABI 3130 Genetic Analyzer according to manufacturer’s protocol.

To generate pCDNA3.1-c.336C>GPAX9-myc-His(A), *in vitro* site directed mutagenesis was performed using QuikChange site directed mutagenesis kit (Stratagene, USA) using manufacturers protocol. In this reaction 5’GGCGGACGGCGTGTG**<u>G</u>**GACAAGTACAATGTG-3’ and 5’-CACATTGTACTTGTC**<u>C</u>**CACACGCCGTCCGCC-3’ were used as mutagenesis template sense and anti-sense respectively (Mutant base is underlined). Introduction of the desired mutation was confirmed by automated DNA sequencing.

### 2.5 Cell culture and transfection

COS7 and HEK293 cells were obtained from National Centre for Cell Science, Pune, India respectively and grown in DMEM (Sigma-Aldrich, USA) supplemented with 10% FBS (Invitrogen, Thermo Fisher Scientific, USA) and 100U/ml of Penicilin-50µg/ml Streptomycin antibiotic (Himedialabs, India) and maintained at 37°C under 5% CO_2_ with >95% relative humidity. At the time of transfection complete DMEM was replaced with Opti-MEM Reduced Serum Medium with GlutaMAX supplement (Invitrogen, Thermo Fisher Scientific, USA) and transfection with plasmid was carried out using Lipofectamine 2000 (Invitrogen, Thermo Fisher Scientific, USA) according to manufacturer’s protocol.

### 2.6 Preparation of PAX9-stable tranfected HEK293 cell line

HEK293 grown in T25 flask was transfected with PAX9-pCMV SPORT6 mammalian expression construct according to previously described protocol. After 72hr incubation transfection medium was replaced with mammalian selection medium consists of DMEM (Dulbecco’s Modified Eagle’s Medium) (Sigma-Aldrich, USA) supplemented with 10% Fetal Bovine Serum (FBS) (Invitrogen, USA), and antibiotic (100U/ml Penicillin, 100μg/ml Streptomycin) (HiMedia, India) and 600μg/ml of G-418 (Thermo Fisher Scientific, USA). Old selection medium was replaced with fresh selection medium in every 2 days and after around 7 days massive cell death was visible. Cell line was cultured for around 20 days with regular replacement with fresh selection medium. When cell colonies were visible, they were treated with 0.1%-Trypsin-0.1% EDTA as described previously and cultured for additional 15 days. After that a fraction of cells were used for DNA isolation according to (Sambrook and Russell, 2001). Briefly, medium was discarded from 80% confluent cells attached to T25 flask and cells were washed with PBS. 1ml DNA extraction buffer was added to it and incubated at 55°C for 8hr. DNA was precipitated by NaCl-Ethanol solution followed by washing of DNA with 70% ethanol. PCR to amplify DNA fragment which can represent the transfected plasmid clone was carried from the isolated genomic DNA using appropriate primer.

### 2.7 Protein expression and subcellular localization

To study the expression of wild type and mutant PAX9 protein, COS7 and HEK293 cells were transfected with wild type [pCDNA3.1-PAX9-*myc*-His(A)] and mutant [pCDNA3.1-c.336C>GPAX9-*myc*-His(A)] *PAX9* construct and total cellular protein was isolated using modified RIPA buffer after 48 hr of transfection. Protein samples were resolved using 10% SDS-PAGE and analyzed by Western blotting using anti-c-Myc antibody (Sigma-Aldrich, USA) and anti-PAX9 antibody (Cell signalling technology, Inc., USA). Empty vector transfected cell lysate was used as negative control. For sub-cellular localization study cell line was fixed using 3.7% paraformaldehyde after 48 hr of transfection. Permeabilization and blocking were carried out for 30 min at room temperature using 0.1% PBST containing 1X PBS, 0.1% sodium deoxycholate (Sigma-Aldrich, USA), 0.1% Triton X-100 (Sigma-Aldrich, USA), 0.1%BSA and 10% FBS.Cells were incubated with anti-c-Myc antibody (Clontech Laboratories, USA) at a 1:10 dilution for 16 hr at 4°C, washed and incubated with incubated with Cy3 conjugated goat anti-mouse antibody (Sigma-Aldrich, USA). Cells were counter stained using DAPI (4’, 6-diamidin0-2-phenylidone dihydrochloride, Sigma-Aldrich, USA) and mounted in DABCO (antifading agent, Sigma-Aldrich, USA). Images were collected using laser scanning confocal microscope LSM510-Meta (Carl Zeiss) and analyzed using LSM510-Meta software. To cross verify immunocytochemistry results, nuclear and cytoplasmic fraction were prepared from 48 hr transfected cells according to Dimauro *et al*. 2012 with slight modifications and was analyzed by Western blotting using anti-c-Myc antibody (Sigma-Aldrich, USA) and anti-PAX9 antibody (Cell signalling technology, Inc., USA).

### 2.8 Electrophoretic Mobility Shift Assay (EMSA) for DNA-Protein interaction study

Putative paired domain binding sequence e5 (39-mer) from *Drosophila even-skipped* promoter, as described previously (Mensah et al., 2004), was used as a probe to study DNA binding activity of mutant PAX9 protein. Sense, (5-CACCGCACGATTAGCACCGTTCCGCTCAGGCTGTCCCCC-3’) andantisense, (5’-CACGGGACAGCCTGAGCGGAACGGTGCTAATCGTGCGGT-3’) strand of e5 probe were synthesized (Sigma-Aldrich) and sense strand was end labelled with [γ^32^P] ATP using T4 polynucleotide kinase according to manufacturer’s protocol. Unincorporated nucleotides were removed by Sephadex G-25 size exclusion chromatography. Equimolar amount of labelled sense strand and unlabelled antisense strand were annealed to generate radio labelled double stranded oligonucleotide probe. Optimum amount of labelled probe used for EMSA experiment was empirically determined by resolving different amounts of labelled probe by Polyacrylamide gel electrophoresis (PAGE) followed by autoradiography. Appropriate amount of labelled probe was incubated with 6µg nuclear lysate from COS7 cell line transfected separately with wild type [pCDNA3.1-PAX9-*myc*-His(A)], mutant [pCDNA3.1-c.336C>GPAX9-*myc*-His(A)] *PAX9* and pCDNA™3.1-*myc*-His(A). Super shift experiment was performed by adding 1 µg anti-c-Myc antibody (Sigma-Aldrich, USA) to each protein-DNA complex followed by 3 hr incubation in ice. Protein-DNA and antibody-protein-DNA complex was resolved by 6% Polyacrylamide gel electrophoresis (PAGE) followed by autoradiography.

### 2.9 Expression of GST tagged protein

GST-fused bacterial expression plasmid for *PAX9* was created by subcloning wild type and mutant (c.336C>G) *PAX9* in BamHI and SalI site of pGEX4T1 vector (GE Healthcare, USA). *E. coli* BL21 competent cells were transformed with each recombinant plasmid. Single colony was inoculated in 5 ml LB broth supplemented with 500 µg of Ampicillin and cultured for overnight at 37°C with shaking in 200 rpm. 100 ml LB broth containing 100 µg/ml Ampicillin was inoculated with 1 ml saturated culture and cultured at 37°C until OD_600_ reaches to 0.45. Incubation temperature was reduced to 16°C and culture was induced with 0.1 mM Isopropyl-1-thio-β-galactopyranosid (IPTG). Cultures were harvested after 10 hr by centrifuging at 5000g for at 4°C for 5 min. and resuspended in 2ml of lysis buffer. Cell lysates were prepared by sonication and centrifuged at 13000g for 30 min at 4°C. Supernatant containing wild type and mutant PAX9 was used for GST pull down as bait protein.

### 2.10 GST-Pull down experiment for protein-protein Interaction study

FLAG-*MSX1* cloned in pCDNA3.1V5-His was expressed in COS7 cell line and was used as prey protein. Total cellular protein from FLAG-*MSX1*-pCDNA3.1V5-His transfected COS7 was isolated using standard protocol. Briefly, transfected cells were washed with chilled PBS and lysed in appropriate lysis buffer (50 mM Tris-Cl pH 8.0, 200 mM NaCl, 5% glycerol,0.5% Triton-X 100, 1 mM PMSF, 1X Protease inhibitor, 1X Phosphostop) after incubating in ice for 1 hr followed by centrifugation at 13000g for 30 min at 4ºC. Supernatant containing FLAG-MSX1 was incubated for 10 hr at 4°C with GST-PAX9 (wild type and mutant) protein bound to glutathione-sepharose resin. Protein complexes were washed six times with buffer containing 50 mM Tris-Cl pH 8.0, 200mM NaCl, 0.5% NP40. Protein bound beads were boiled with 1X SDS loading buffer and resolved in 10% SDS-PAGE and analyzed by with 1:1000 dilution of rabbit anti-PAX9 polyclonal antibody (Cell Signaling Technology, USA) or 1:1000 dilution of mouse anti-FLAGM2 monoclonal antibody (Sigma-Aldrich, USA).

### 2.11 RNA isolation from cell line and cDNA synthesis

Total cellular RNA was isolated from cell line using TRI reagent (Sigma-Aldrich, USA). Briefly, culture medium was removed from 90% confluent cells and washed with PBS. To the cells appropriate volume of TRI reagent was added and then cell lysate was passed several times through pipette and transferred to a sterile 1.5ml micro centrifuge tube. To the lysate, 0.2ml of chloroform was added per 1ml of TRI reagent, mixed by vortexing and kept at room temperature for 15 min followed by centrifugation at 12000g for 15min. at 4°C. Upper aqueous phase was collected in a fresh tube and 0.5ml of 2-proponal was added per 1ml of TRI reagent, mixed by vortexing and kept at room temperature for 10 min. followed be centrifugation at 12000g for 10min. at 4°C. RNA pellet was washed with 75% ethanol, air dried and dissolved in 20μl of nuclease free water. RNA was quantified in spectrophotometer and checked by 1% agarose gel stained with EtBr. RNA was treated with DNase I (RNase free) (Ambion) according to manufacturers’ protocol. Briefly, a 30μl reaction volume containing 30μg of total cellular RNA, 1X reaction buffer, 6U of DNase I (RNase free) and nuclease free water. Reaction was incubated at 37°C for 30min. After incubation 70μl DEPC water was added to the reaction and RNA was purified by 100μl TRI reagent as described previously.

### 2.12 cDNA synthesis and Real time PCR

Random priming was used to synthesise cDNA from 2μg of total cellular RNA using High-Capacity cDNA Reverse Transcription Kit (Applied Biosystems, USA) according to manufacturers’ protocol. cDNA was diluted 10 times with autoclaved deionised water. 5μl of diluted cDNA was used for Real-Time PCR using appropriate primers and SYBR Green Real-Time PCR Master Mix (Thermo Fisher Scientific, USA) in ABI 7500 Real-Time PCR System (Applied Biosystems, USA)

### 2.13 RNA sequencing

Total cellular RNA was used for library preparation using TrueSeq RNA sample preparation kit V2 (Illumina). Library QC was performed using Agilent Bioanalyzer 2100 followed by quantification by real time PCR using KAPA NGS library quantification kit (KAPA biosystem). Cluster was generated in cBot instrument (Illumina). Sequencing by synthesis was carried out in Illumina HiSeq 2000 platform for 50bp single end read.

Base call files (.bcl) generated by Illumina HiSeq 2000 platform were converted into nucleotide sequence with quality score FASTQ (FASTA with quality score) using CASAVA software v1.8.2 (Illumina) followed by adaptor trimming. Bases with QC<30 were discarded. Filtered reads were aligned to human genome and transcriptome data hg19 (UCSC) build followed by normalization and quantification using DEseq algorithm available with Avadis NGS v1.5.1 (Strand scientific intelligence, Inc). Differentially expressed gene list was analysed for pathway analysis using Avadis NGS v1.5.1 (Strand scientific intelligence, Inc).

## 3 Results

### 3.1 Clinical findings and inheritance pattern

In DEN14 family, information regarding congenital tooth agenesis was traced back to four generations. In total 12 individuals, including 10 females and 2 males, were reported to be affected with tooth agenesis in this family. The proband (IV7), an 8 years old girl, was affected with autosomal dominant type of posterior tooth agenesis, which includes 4 mandibular molars, 2 maxillary molars and 2 maxillary canines (Fig.1 A & B). In the proband status of third molars was not determined due to lack of calcification of tooth bud at this age. Out of total permanent missing teeth, contribution of molars (excluding third molars) was highest (89.6%) followed by premolars (6.3%), incisors (4.2%) and canines (4.2%). Dental status for other patients in the generation III appeared similar but could not be verified for individual teeth because of the lack of orthopantomograms (Fig. 1C).

**Fig1.**
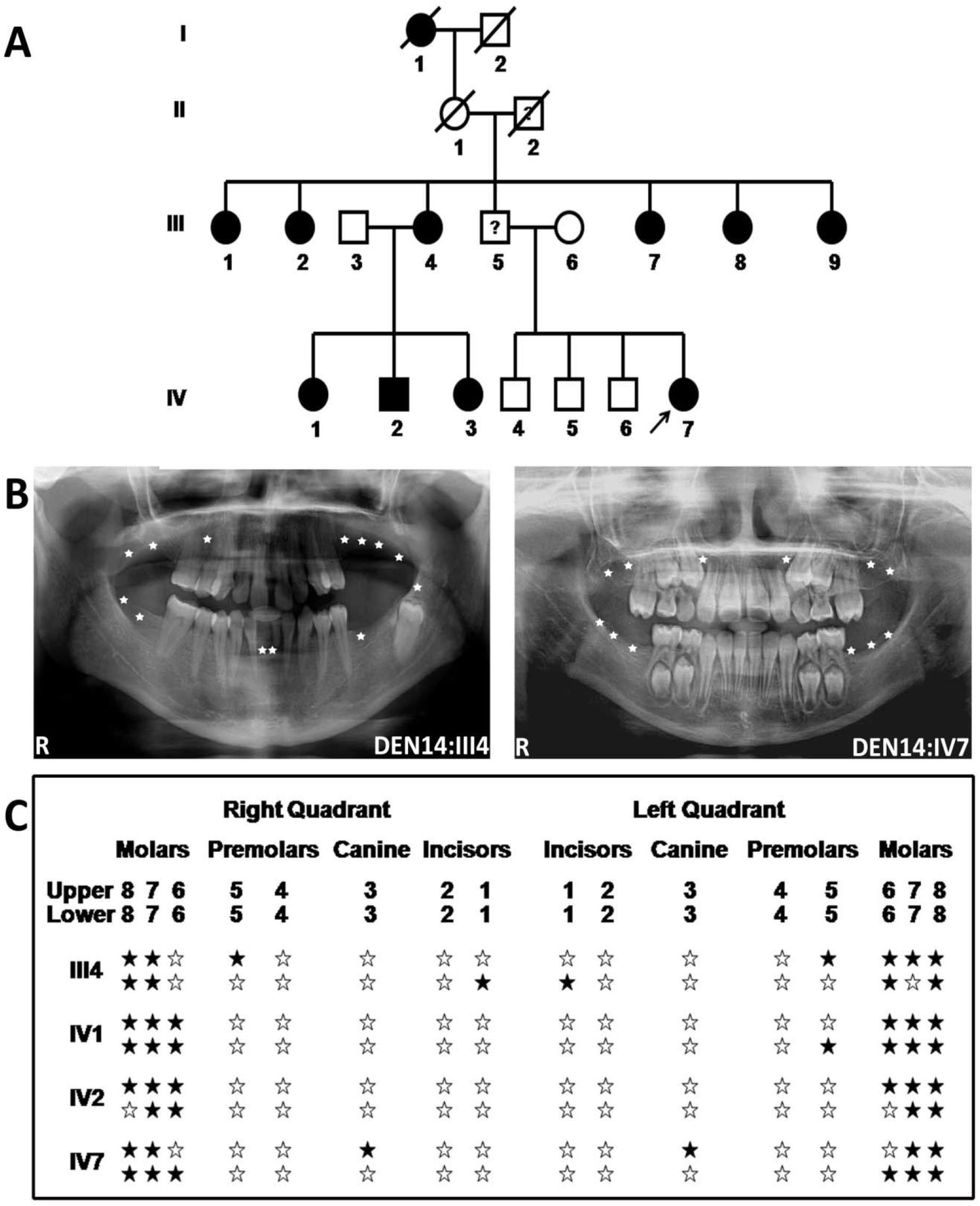
Pedigree structure and tooth agenesis pattern in affected individuals of DEN14 family. (A) Pedigree showing pattern of inheritance of congenital tooth agenesis in DEN14 family. (*Squares* males, *circles* females, *blackened* affected, *clear* unaffected, *arrow* proband, *slash* deceased). (B) Representative orthopantomogram (OPG) of the proband showing the tooth agenesis pattern (*Star* missing permanent teeth). (C) Permanent dentition pattern of representative affected individuals. (*Star* position of permanent teeth, *filled star* missing permanent teeth.

There was no report of deciduous tooth agenesis in this DEN 14 family. No clinical abnormality related to hair, nail, skin and sweat gland, sweating activity as well as mental retardation or cancer was reported.

### 3.2 Identification of novel c.336C>G mutation

Direct sequencing of all the coding exons and exon-intron boundaries of *PAX9* and *MSX1* along with putative promoter region of *PAX9* identify a potentially novel cytosine to guanine transversion at c.336 position of *PAX9* gene (Fig. 2). All the affected individuals were detected to carry this variation in heterozygous condition but two studied unaffected family members did not carry this variation. No novel or known pathogenic mutation was observed in *MSX1*. This variation was not reported in NCBI-dbSNP, ClinVar and 1000 Genome data base. None of the 200 control chromosomes from same ethnic background carried this variation. This novel was submitted to NCBI ClinVar database (Accession Number. VCV000155939.1; https://preview.ncbi.nlm.nih.gov/clinvar/variation/155939/evidence/)

**Fig2.**
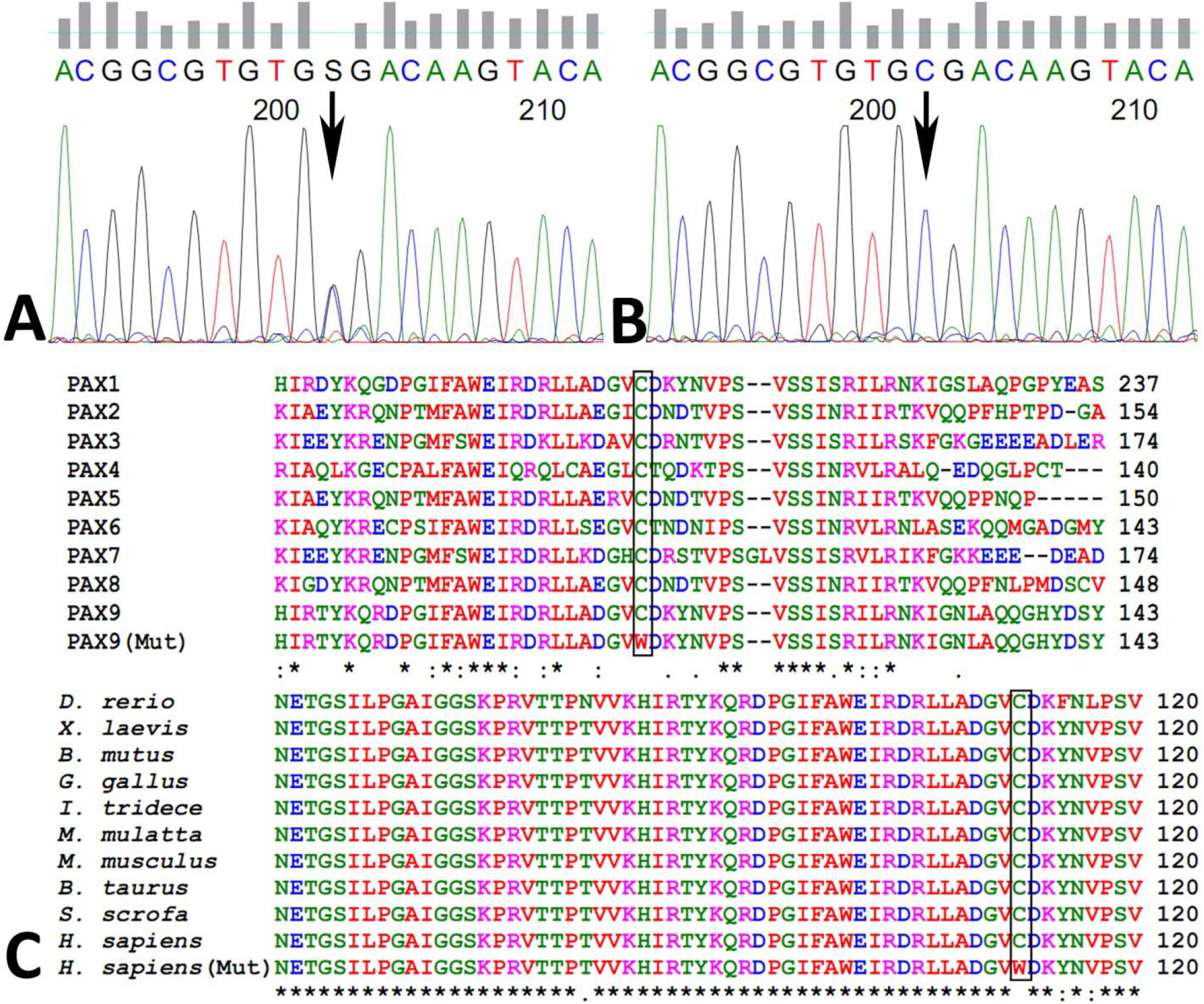
Identification of novel c.336C>G variation in *PAX9*. Representative DNA chromatograms showing (A) wild-type homozygous ‘C’ at c.336 position in unaffected individuals and (B) heterozygous CG in affected individuals in second coding exon (Exon 3) of *PAX9*. (C) Multiple sequence analysis of human PAX1 to PAX9 protein (Upper) and multiple sequence analysis of PAX9 across different vertebrate species (Lower). This showing conserved Cys residues (within box) across different PAX family members and vertebrate species.

### 3.3 In silico analysis of PAX9 (p.Cys112Trp) variation

Multiple sequence analysis of PAX9 across different vertebrate species and all the nine members of human PAX proteins using Clustal W2 software, showed that Cys112 residue of PAX9 is evolutionary conserved (Fig. 3). Subsequent analysis of p.Cys112Trp variation using PolyPhen-2 and MutationTaster predicted them as ‘probably damaging’ or ‘predicted disease causing’.

**Fig3.**
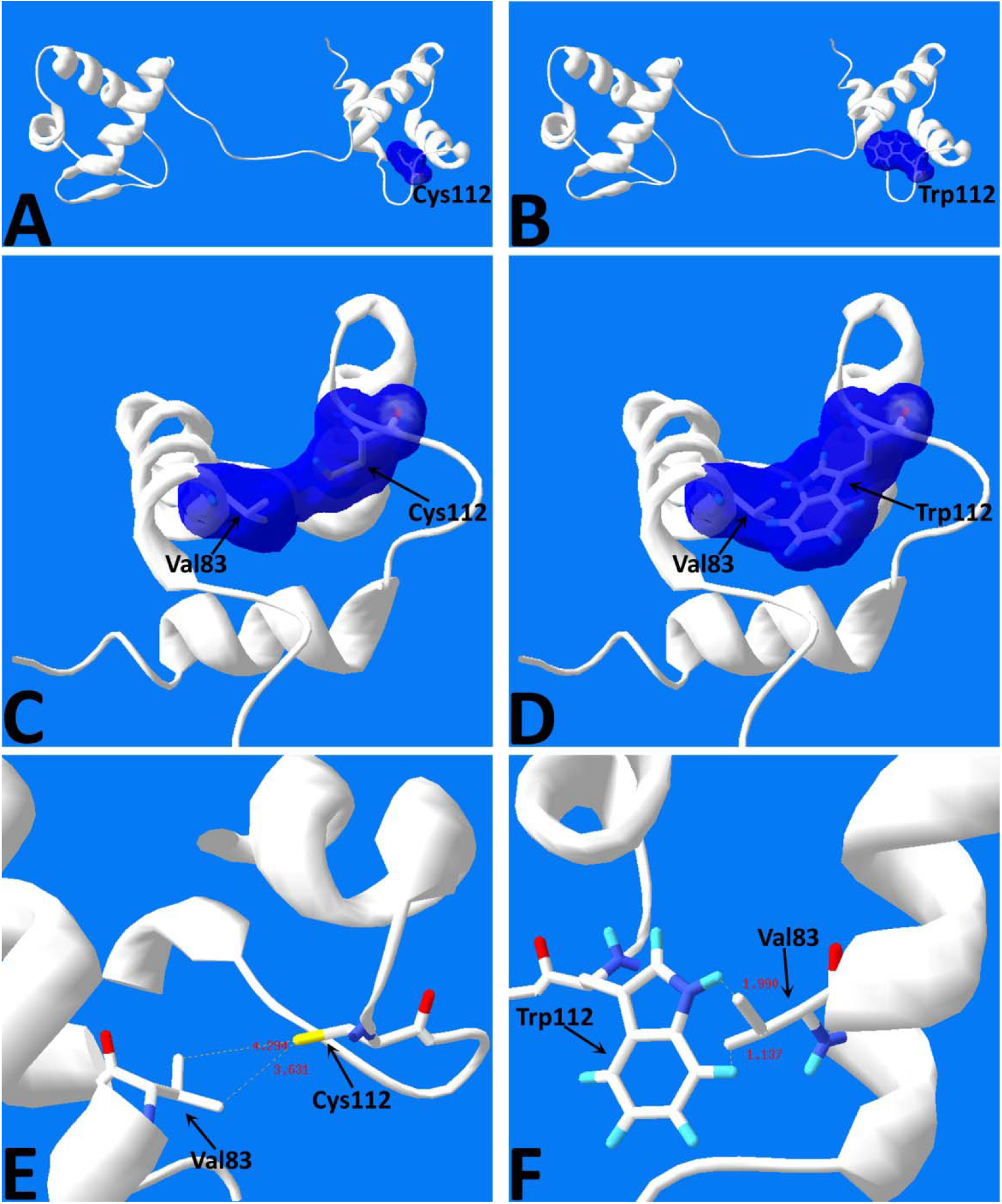
*In silico* analysis of p.Cys319Trp mutation on protein structure of PAX9-DNA-binding paired domain. (A) location of Cys112 at paired domain region of wtPAX9 (B) location of Trp112 in mutPAX9 (C) location and surface overlapping of Val83 and Cys112 (D) location and surface overlapping of Val83 and Trp112 (E & F) distance between closest atom of Cys112 and Val83 and that of Trp112 and Val83.

The structure of wild and mutant proteins based on homology modelling has more than 90% residues in the most preferred region. That showed a good model (Fig.4). Superposition of wild and mutant PAX9 showed structural anomalies noted in mutant PAX9 (Fig 5). Template based homology modelling indicated that the Cys112 is located within the linker region between α-helix 2 and α-helix 3 of C-terminal sub-domain of the DNA-binding domain of PAX9. Secondary structure prediction showed changes in helix, putative domain and boundary coil regions in mutant PAX9 (Fig. 6).

**Fig4.**
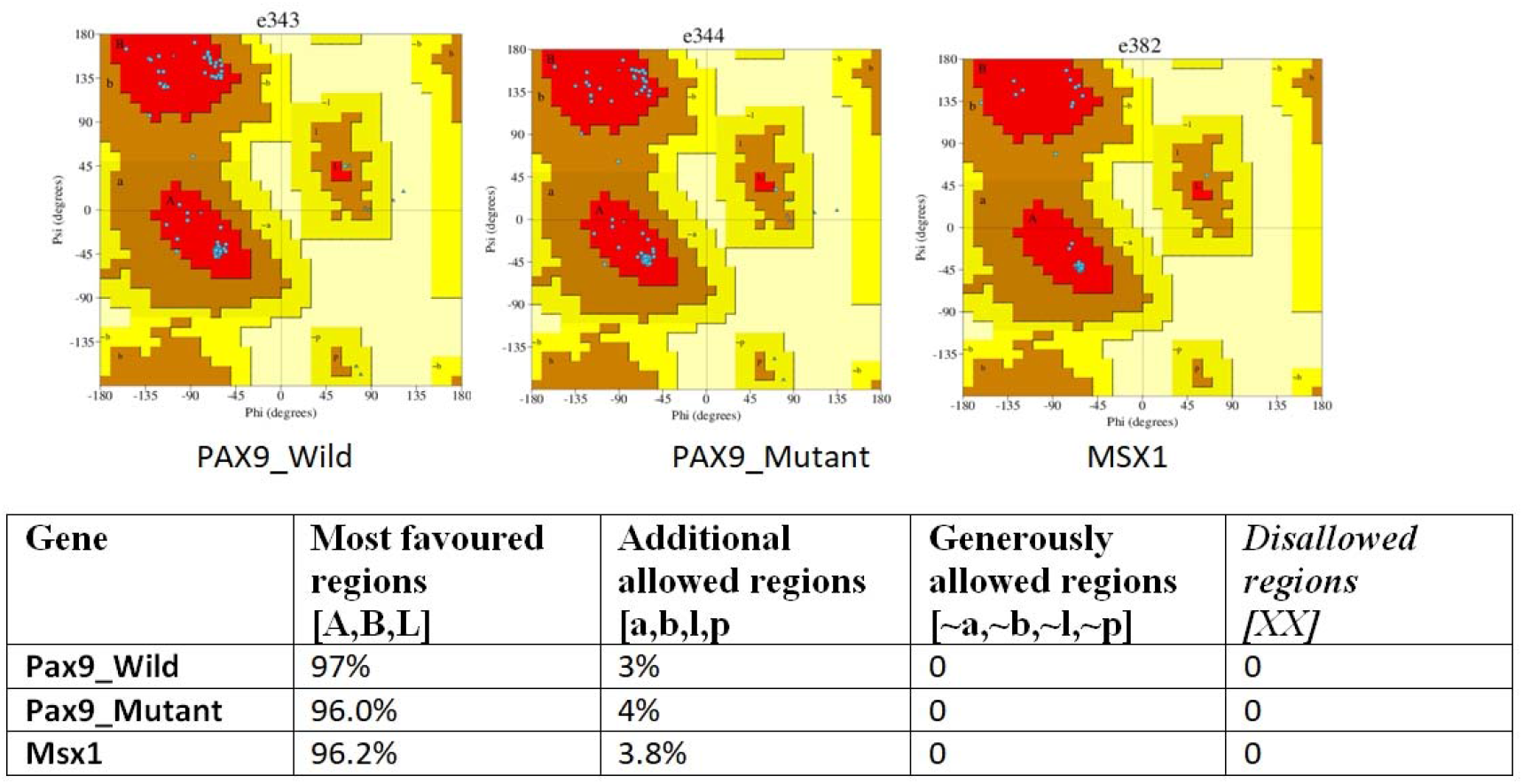
Evaluation of protein structure by Ramachandran plot analysis.

**Fig5.**
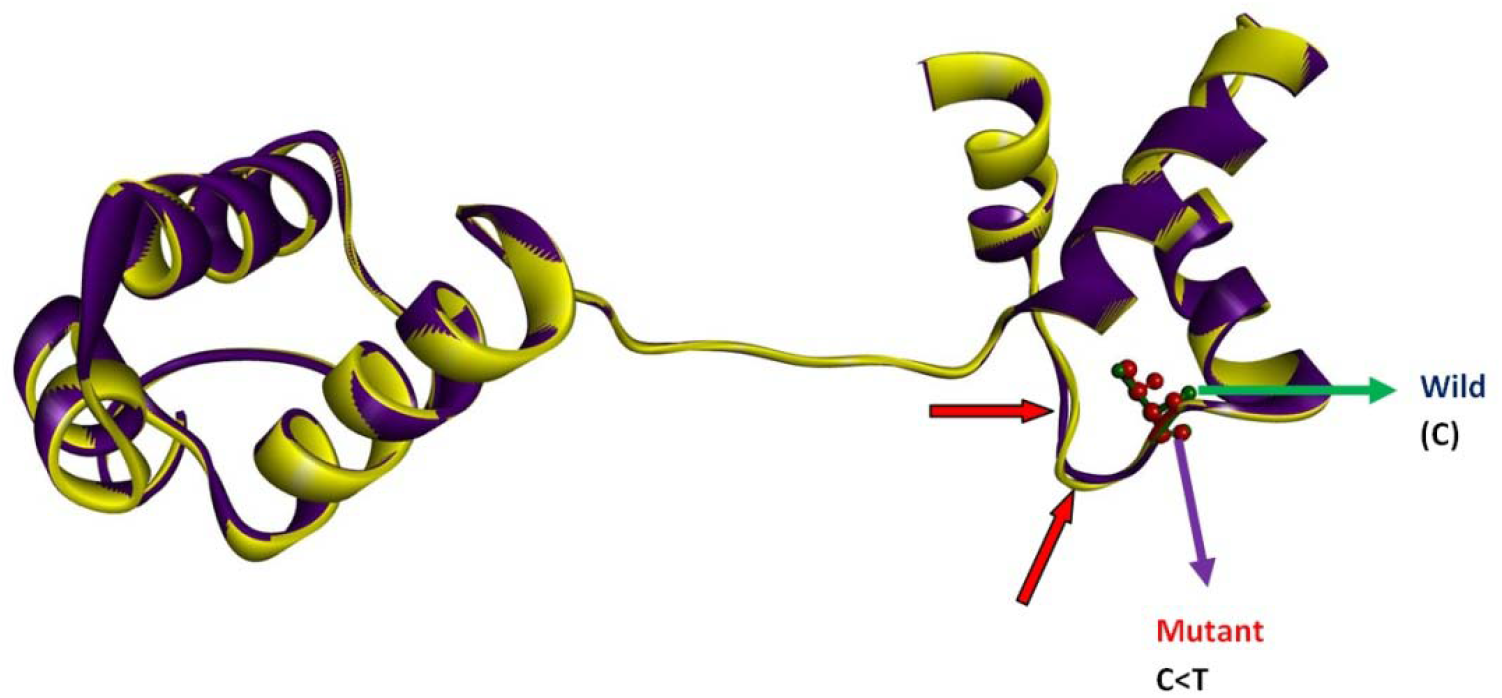
Superposition of wild (Blue) and mutant (Yellow) PAX9. Red arrow indicates structure conformation hampered.

**Fig6.**
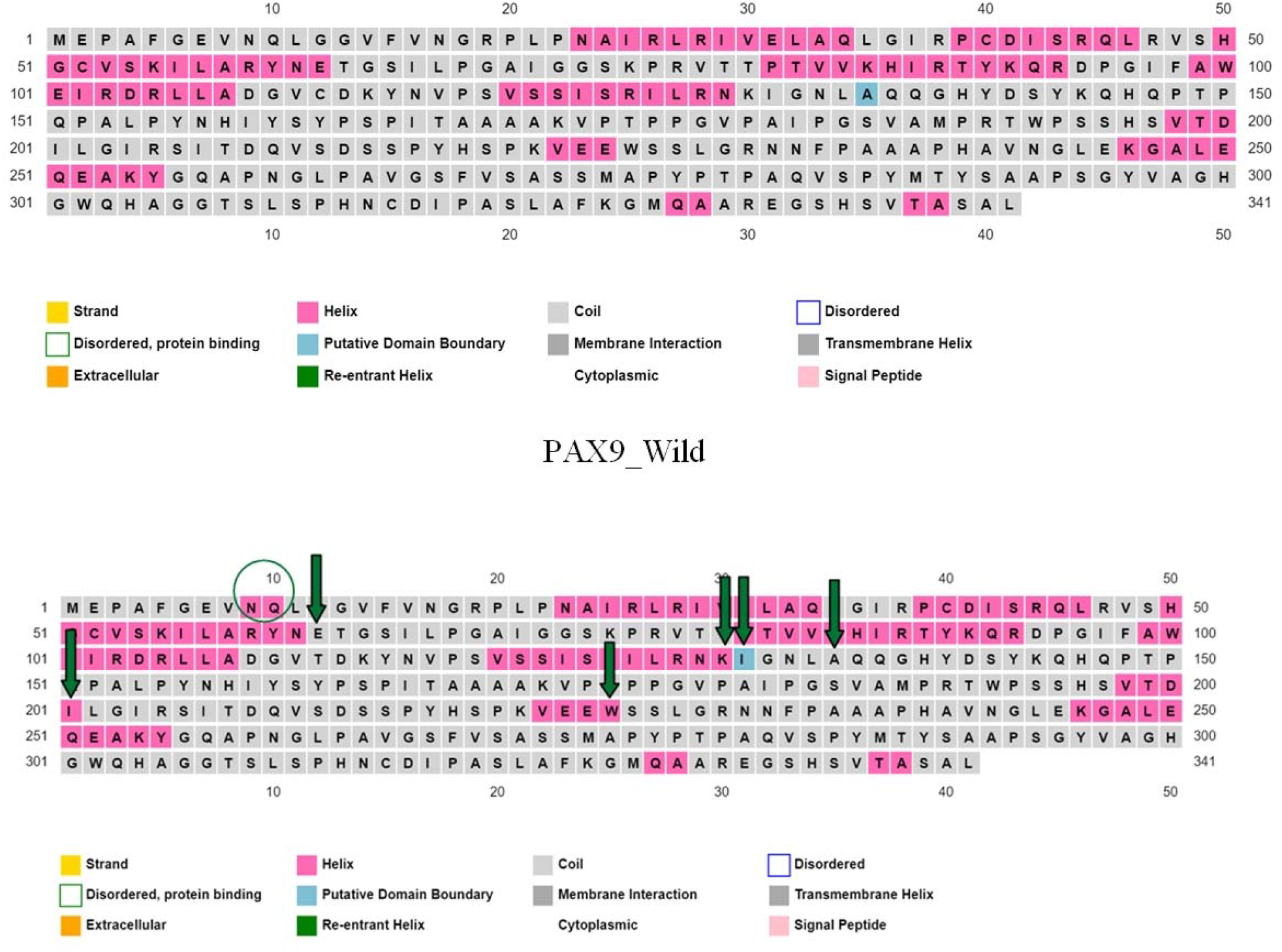
Secondary structure prediction of Wild and mutant of PAX9.

Docking study reveals that there is significant difference in the number of salt bridges, hydrogen bonds, disulphide bonds and non-bonds contacts between wild type and mutant (Fig.7) PAX9 while they interact with MSX. This leads to reduce the binding affinity of mutant PAX9 with MSX1 compared to wild type PAX9 (Table1) (Fig.8). DNA-Protein interaction study using PatchDock indicates reduced binding affinity of mutant PAX9 with e5 probe. The wild complex had a docking score of 14870, while the mutant complex had a score of 14652 (Fig9). MDS study showed that mutant PAX9 has lower RMSD value compared to wild type PAX9, which predict that the mutant protein will be more stable (Fig10.). Radius of gyration (RG) value of mutant protein is lower than wild type making the mutant protein more rigid (Fig10.). Along with that mutant protein shows less solvent accessible surface. (Fig10.).

**Table1:** Binding energy and other parameters comparisons between wild and mutant complex.

|  | PAX9<sub>wild</sub>-MSX1 | PAX9<sub>mut</sub>-MSX1 |
| --- | --- | --- |
| Binding Energy(Kcal/mol) | -13.9 | -10.2 |
| No. of intermolecular contacts | 108 | 80 |
| No. of interfacial contacts(ICs) per property |  |  |
| ICs charged-charged | 12 | 8 |
| ICs charged-polar | 18 | 10 |
| ICs charged-apolar | 31 | 24 |
| ICs polar-polar | 2 | 6 |
| ICs polar-apolar | 26 | 17 |
| ICs apolar-apolar | 19 | 15 |
| Non interacting surface (NIS) per property |  |  |
| NIS Charged | 43.23% | 41.18% |
| NIS apolar | 27.10% | 28.10 % |
| No. of salt bridges | 1 | - |
| No. of hydrogen bonds | 4 | 8 |
| No. of non-bonded contacts | 249 | 385 |

**Fig 7.**
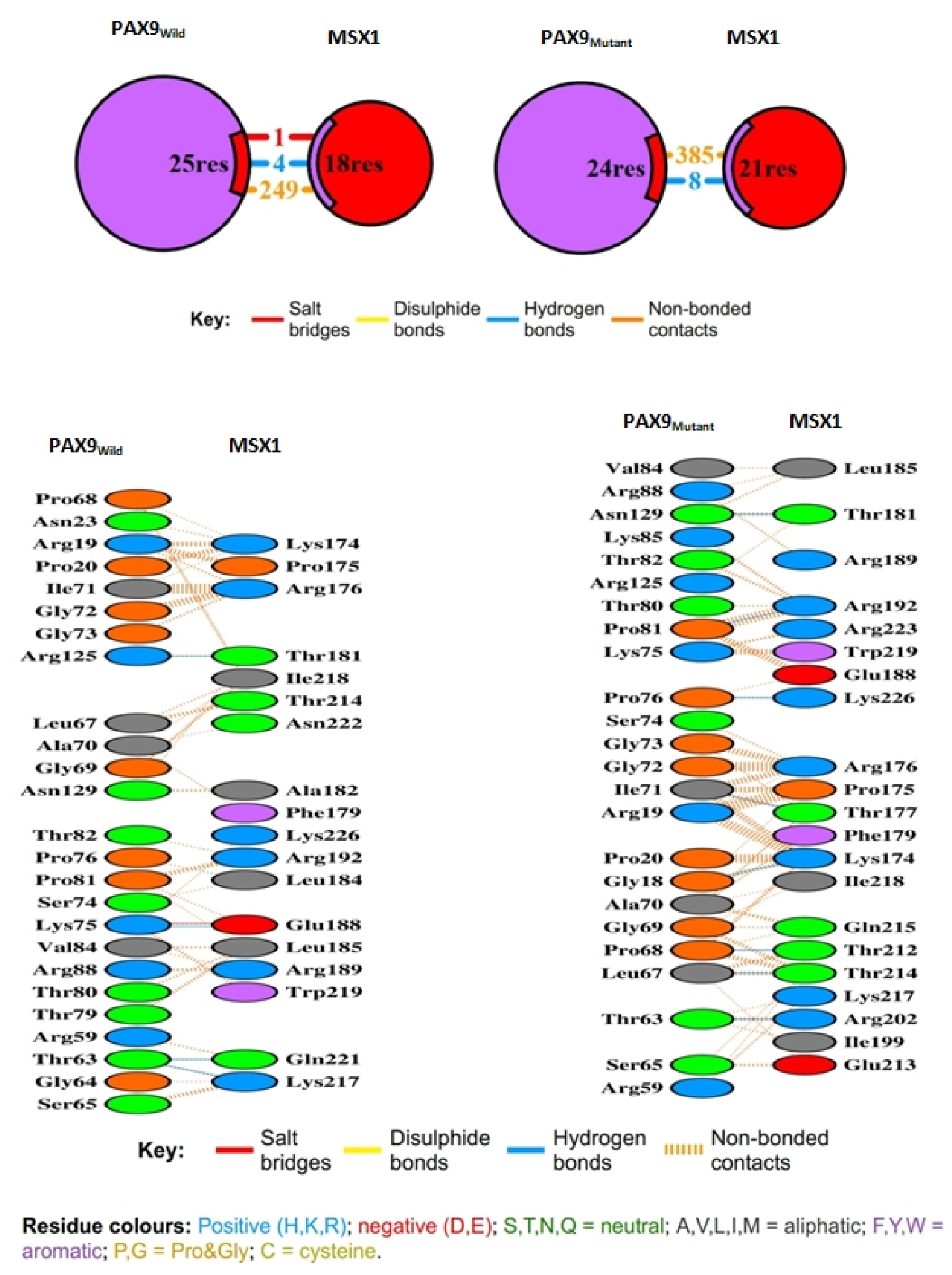
Visualization of interaction between wild and mutant PAX9 with MSX1.

**Fig8.**
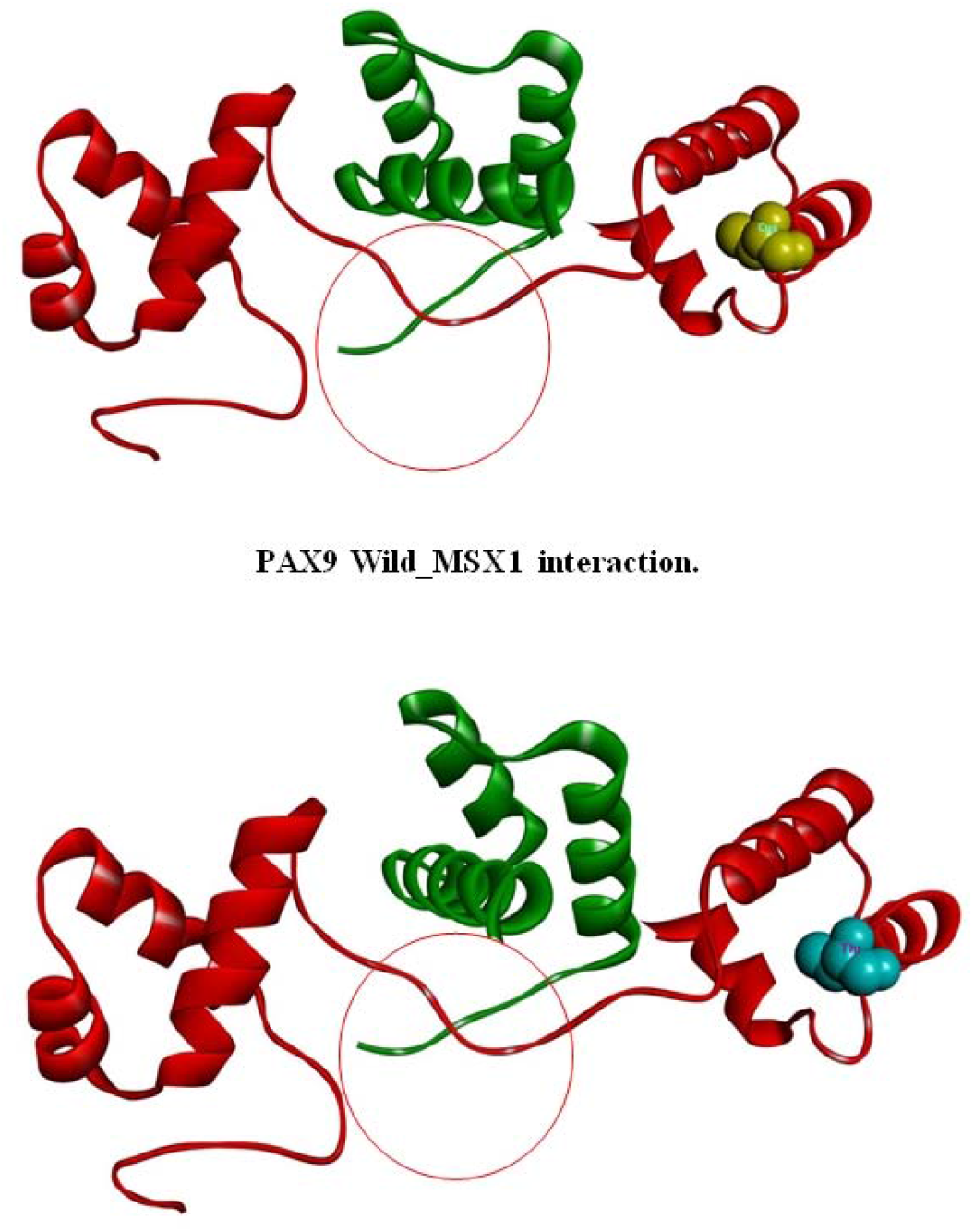
Structural visualization of protein complex of PAX9 wild and mutant with MSX1.

**Fig9.**
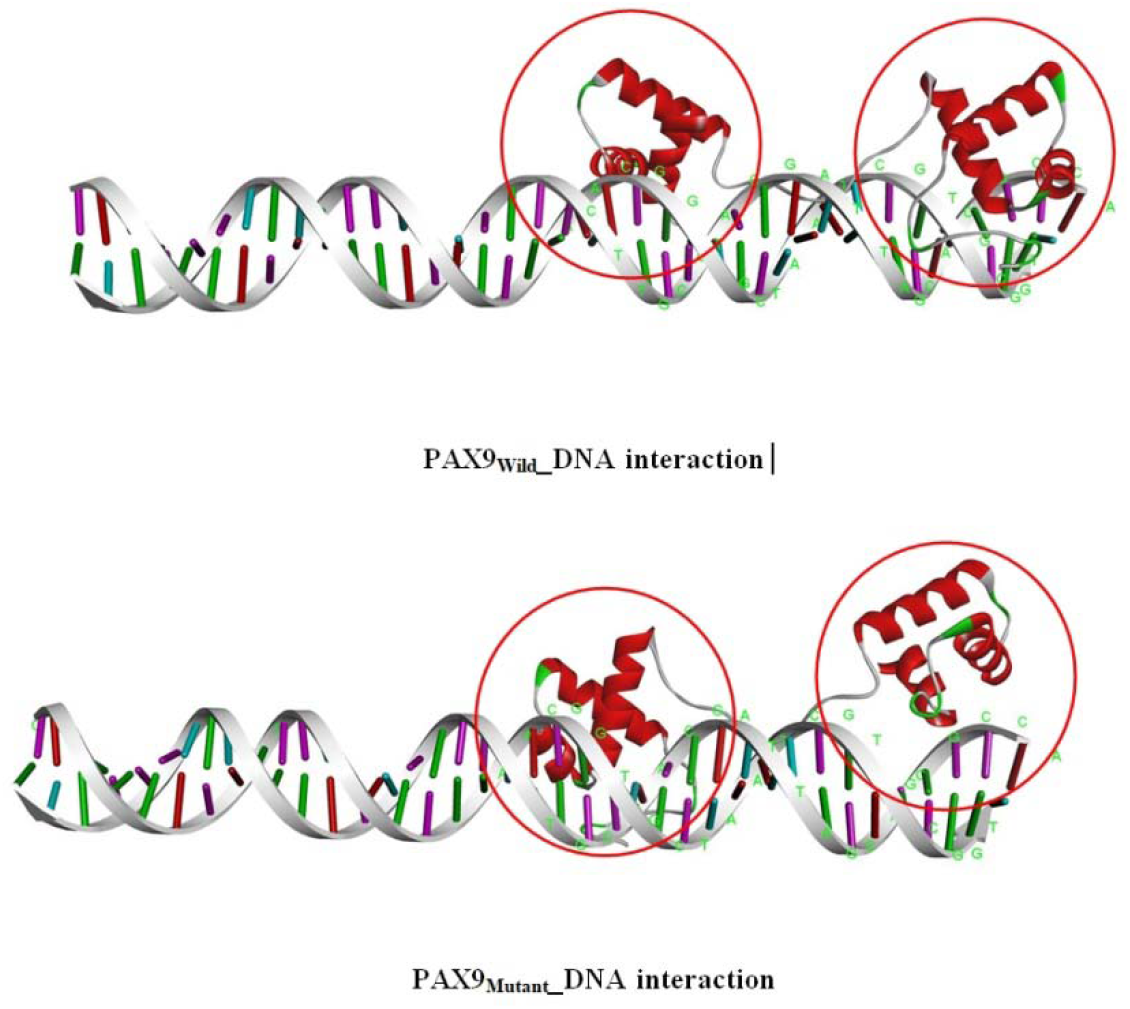
Visualization of the protein-DNA complex structurally (PAX9 wild and mutant with DNA). The red circle denotes the changes that occur in certain areas as a result of contact.

**Fig10.**
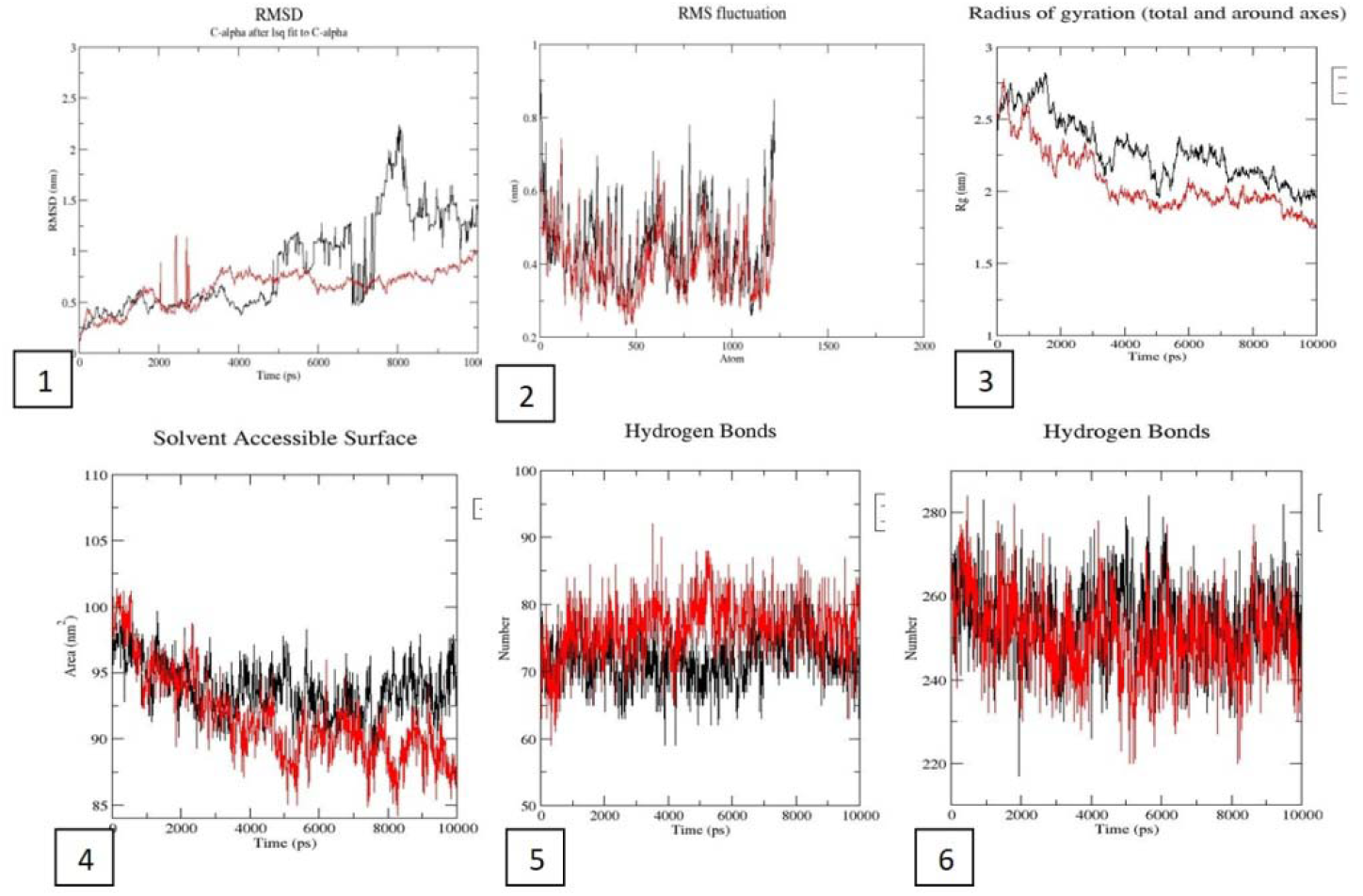
Comparative analysis of MD simulation results of PAX9 wild and mutant. 1.RMSD 2. RMSF 3. Radius of gyration 4. SASA 5. Intramolecular hydrogen bonding and 6. Hydrogen bonding between Protein and water solvent. Black peak indicates wild and red showed mutant.

### 3.4 Nuclear localization unaltered but cytoplasmic retention increased by C112W

Immonocytochemistry result shows both wild type and mutant proteins were predominantly localized at nucleus at 48 hr post transfection in COS7 cell line. However, for mutant protein a higher cytoplasmic retention was observed in17 % of cells transfected with a mutated construct but was observed in only 4 % of cells transfected with the wild type PAX9 (Fig. 11). This result was further cross validated by nuclear and cytoplasmic fraction analysis by Western blotting (Fig.12).

**Fig11.**
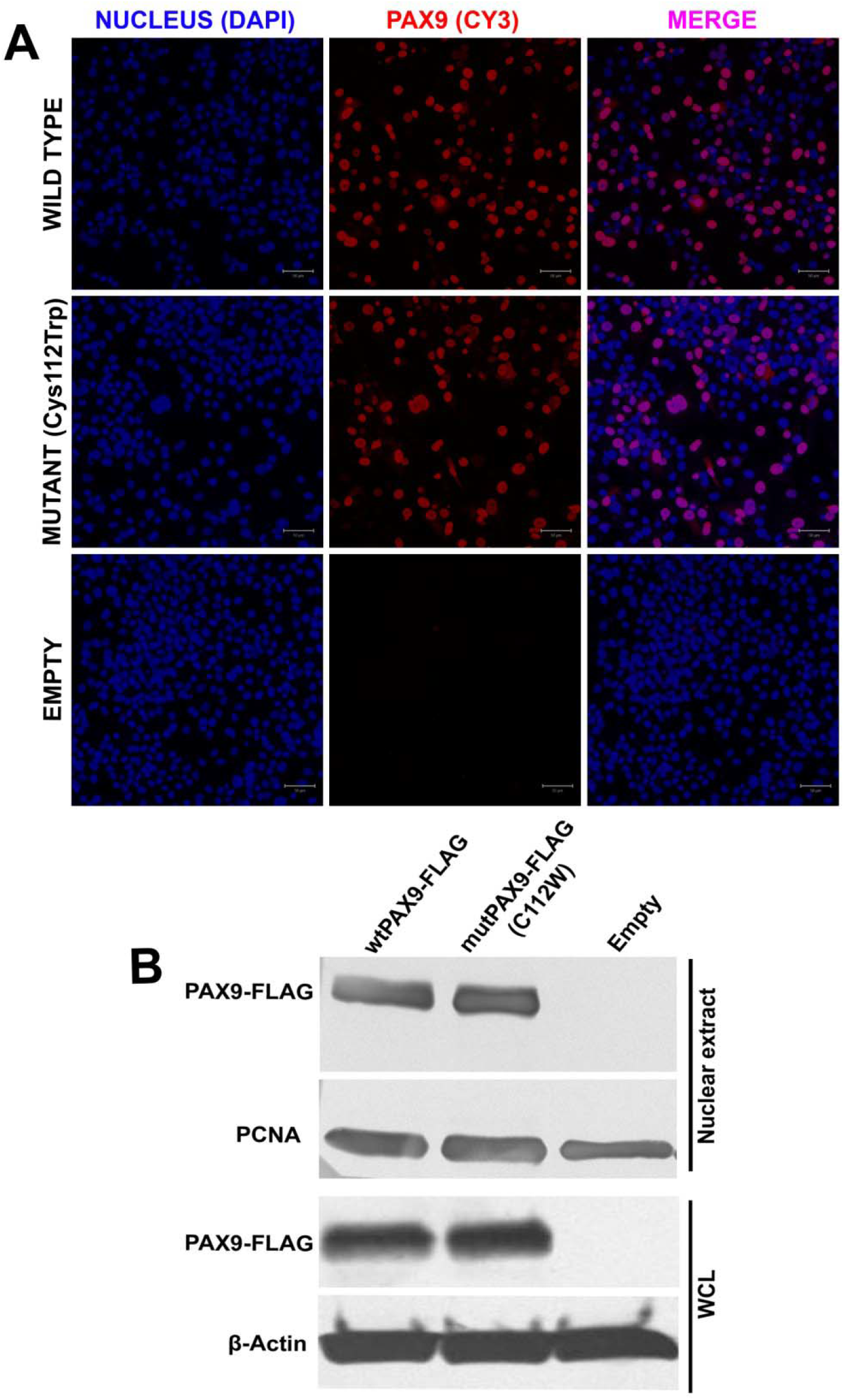
Immuno cytochemistry showing the subcellular localization of wild and mutant (Cys112Trp) PAX9. **A**. Both wild-type and mutant (Cys112Trp) proteins show nuclear localization but the mutant protein shows comparatively higher cytoplasmic retention. **B**. Western blotting showing cytoplasmic and nuclear fraction of wild-type and mutant (Cys112Trp) PAX9 protein. Lane 1 and 2 represents cytoplasmic fraction of 1µg and 0.5μg wt*PAX9*-pCDNA3.1Myc-His transfected cell lysate; lane 6 and 7 represents nuclear fraction of 1µg and 0.5μg wt*PAX9*-pCDNA3.1Myc-His transfected cell lysate; Lane 3 and 4 represents cytoplasmic fraction of 1µg and 0.5μg mut*PAX9*-pCDNA3.1Myc-His transfected cell lysate; lane 8 and 9 represents nuclear fraction of 1µg and 0.5μg mut*PAX9*-pCDNA3.1Myc-His transfected cell lysate; Lane 5 and 10 represent cytoplasmic and nuclear fraction of pCDNA3.1Myc-His transfected cell lysate respectively.

**Fig12.**
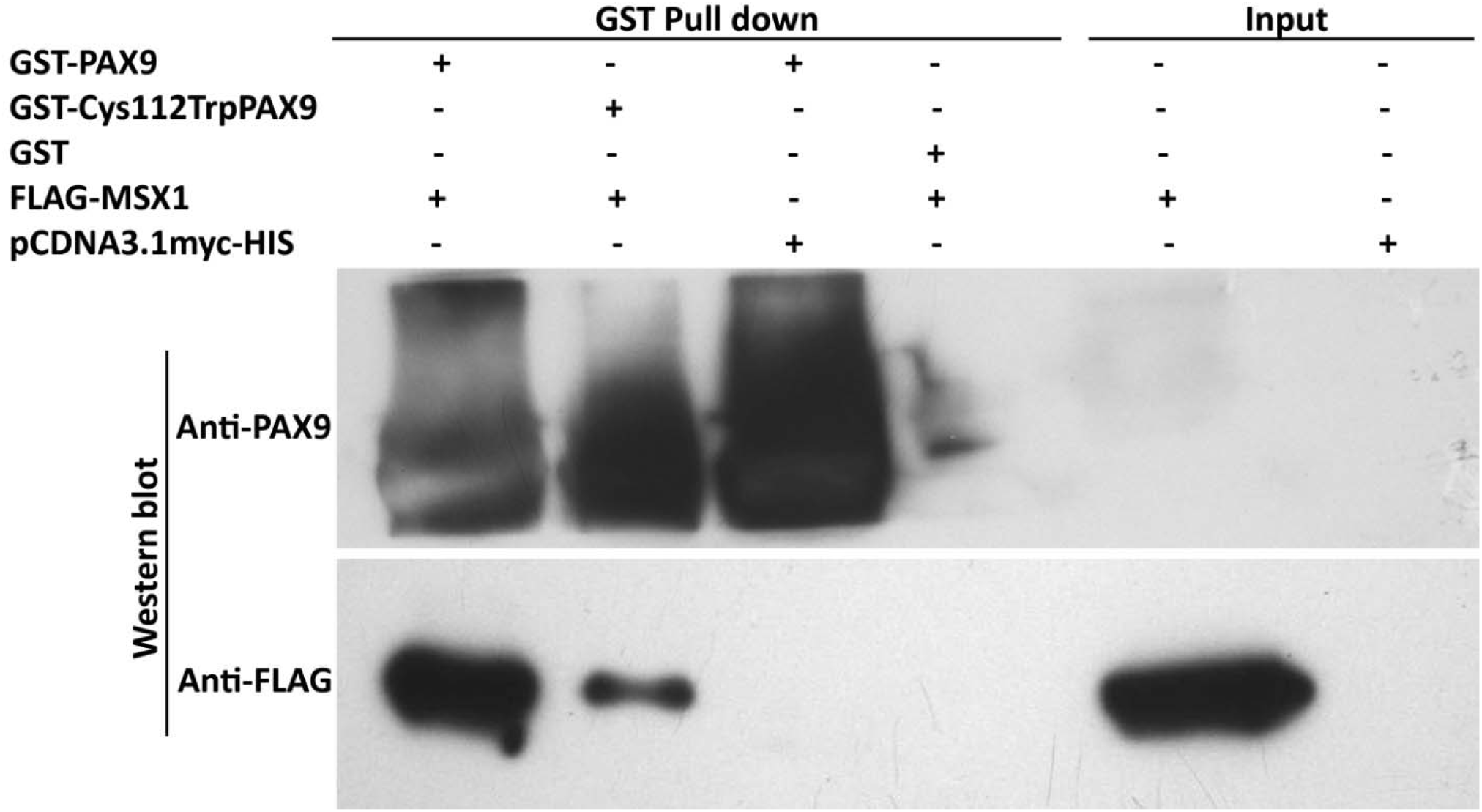
GST pull down assay for protein-protein interaction between MSX1 and PAX9 protein. Western blot of GST pull down assay showing strong interaction between wtPAX9-GST and FLAG-wtMSX1 protein but reduced interaction between Mut(Cys112Trp)PAX9-GST and FLAG-wtMSX1.

### 3.5 C112W mutation reduces affinity for putative paired domain binding DNA sequence

Since Cys to Trp change was located within loop between two C-terminal alpha helices of the conserved paired domain region and it was predicted to have deleterious effect on paired domain structure, sequence specific DNA binding activity of mutant protein was studied through EMSA using radiolabelled *Drosophila* e5 sequence (39-mer) probe. Nuclear lysate from COS7 cell line transfected with wild type *PAX9* clone produce a distinct shifted band of wtPAX9-e5 probe but for mutant *PAX9* no distinct band was observed. Specificity regarding Myc tagged PAX9 binding with probe was verified through super shift assay using anti-cMyc antibody which further shifted up the DNA-protein complex. Thus under *in vitro* condition Cys112Trp mutation disrupt DNA binding activity of PAX9 protein (Fig. 13).

**Fig13.**
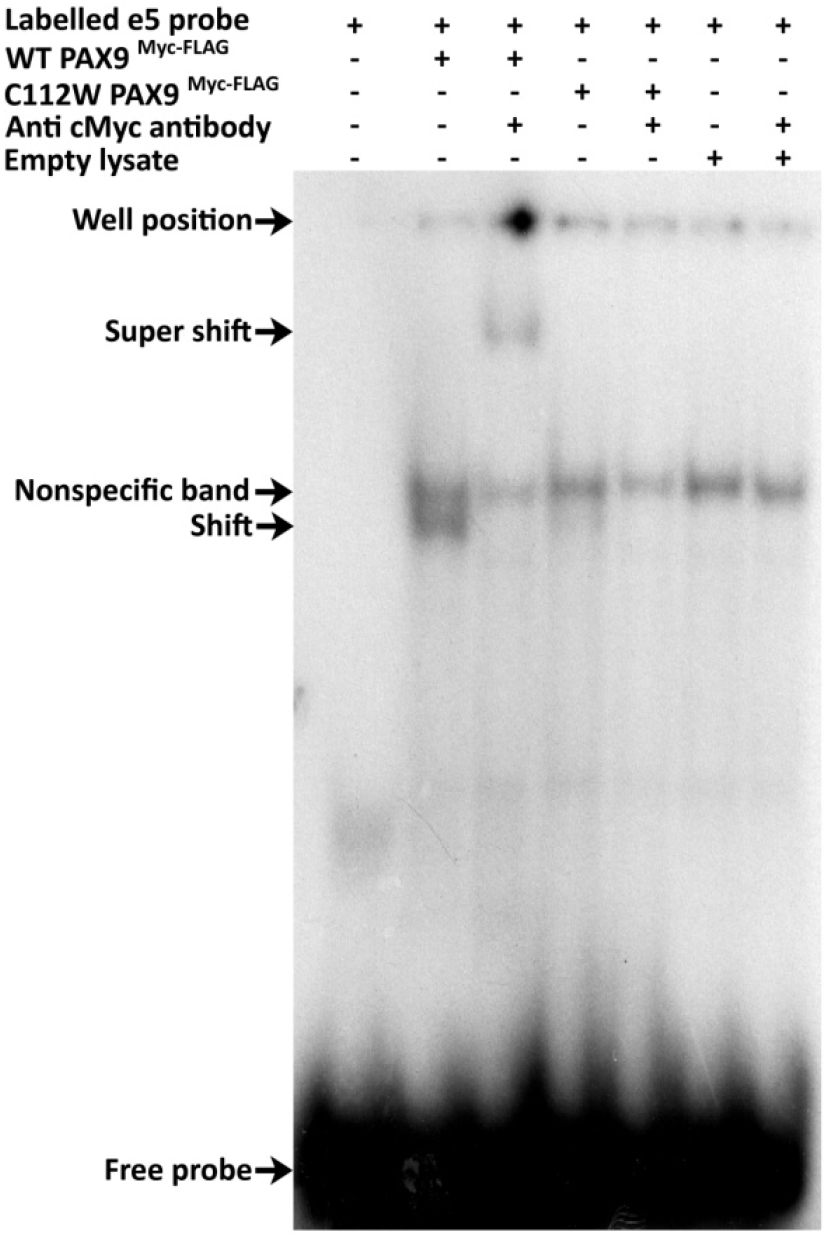
Electrophoretic mobility shift assay to identify DNA binding affinity of wild and mutant (Cys112Trp) PAX9 protein. In the presence of wtPAX9^Myc-FLAG^ a shifted band of e5-PAX9 ^Myc-FLAG^ complex was observed (lane 2) which was further shifted up (super shift) in the presence of anti-Myc antibody (lane 3). Shift and super shift band was not observed in any of the lanes having mutant PAX9 ^Myc-FLAG^ (lane 4 & 5) and also in the control lanes (lane 4 & 5).

### 3.6 Mutant PAX9 (p.Cys112Trp) showed reduced binding with MSX1 protein

Earlier studies in mouse showed that interaction between Pax9 and Msx1 is essential for the stimulation of Bmp4 expression during the bud stage of tooth development. Pax9 are able to interact with Msx1 both in vivo and in vitro (Ogawa et al., 2005; Ogawa et al., 2006). To study the effect of p.Cys112Trp change on the interaction of mutant PAX9 with MSX1 protein GST-Pull down assay was carried out. This experiment showed that Wild-type PAX9 protein bind with MSX1 protein with high affinity but for p.Cys112Trp mutant PAX9 the binding affinity for MSX1 is quite low (Fig. 12).

### 3.7 RNA sequencing and identification of novel PAX9 targeted

To investigate PAX9 regulated genes in mammal, PAX9 stably transfected HEK293 cell line was used as a model. Total cellular RNA was isolated from normal HEK293 and HEK293 cells containing stably transfected PAX9 followed by whole transcriptome analysis through RNA sequencing using Illumina 2500 Next Generation Sequencing platform. Transcriptome profile of these two cell lines were compared using Avadis NGS v.1.5.1 software (Strand Scientific Intelligence, Inc). Compared to untransfected HEK293 cells (Control), 3621 protein-coding genes were differentially regulated in PAX9 transfected HEK293 cells. Out of these 3621 genes, 1730 (47.8%) genes were up-regulated and 1891 (52.2%) genes were down-regulated.

Pathway analysis for differentially expressed genes was performed using pathway analysis module of Avadis NGS v.1.5.1 software (Strand Scientific Intelligence, Inc). 75 pathways were found to be significantly (p<0.05) altered. Among those altered pathways Wnt pathway was well studied during mammalian tooth development and some Pax genes were known to be directly associated with the regulation of Wnt genes (Kim *et al*., 2001, Torban *et al*., 2006). Two WNT pathway genes *WNT4* and *WNT7B* were used for further analysis. For this HEK293 cell line was transiently transfected with wild-type *PAX9* and mutant *PAX9* (NM_006194.3:c.336C>G) and expression level of *WNT4* and *WNT7B* were examined through real time PCR (Fig. 14). Both of the genes were over-expressed in *PAX9* transfected cell line as compared to empty vector transfected and mutant *PAX9* transfected cell line.

**Fig14.**
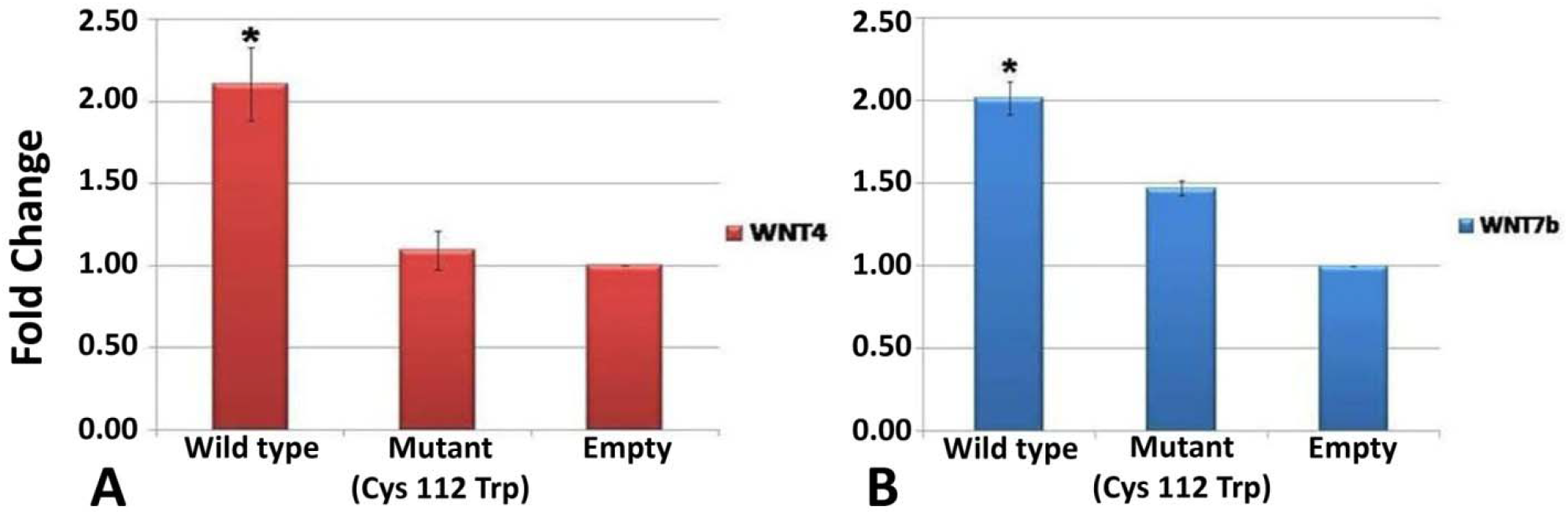
Regulation of *WNT4* and *WNT7b* expression by wild type *PAX9* and mutant *PAX9*. Real time PCR validation of differentially expressed *WNT4* (A) and *WNT7b* (B) gene in wild type *PAX9* and mutant *PAX9* overexpression in HEK293 cells. Data represent mean (±SE) fold change of gene expression. (*p<0.05, two-tail t-Test, equal variance, N=2).

## 4 Discussion

In DEN14 family tooth agenesis was mainly restricted to molars and mutation analysis of PAX9 and MSX1 gene identify novel c.336C>G variation located with the paired domain coding region of PAX9. This variation substitute amino acid Cys with Trp at the amino acid position p.112 (p.Cys112Trp). Cysteine is a small (molecular weight of 121.154g/mol), sulphur containing, polar, uncharged amino acid and on the other hand Tryptophan (molecular weight of 204.229g/mol) is one of the largest non-polar amino acid with bulky aromatic group associated with it.

p.Cys112Trp substitution occurs within a very conserved DNA binding region of PAX9 protein. Multiple sequence alignment shows that Cys112 amino acid is conserved across different vertebrate species and across different members of PAX proteins in human (Fig. 3). From the X-Ray crystallography structures of Pax6-DNA complex and Pax5-DNA complex it was observed that DNA binding domain or Paired domain of PAX protein is consists of 2distint helix-turn-helix (HTH) motif joined by a linker chain. Each of these two HTH motifs consists of 3 α helices joined by short loops which keep those α helices in such an orientation that they can directly interact with DNA (Xu et al., 1999; Garvie et al., 2001). Initial studies regarding DNA binding activity of paired domain using the co-crystal structure of Drosophila Paired-DNA complex (Prd-DNA) was unable to completely determine the DNA binding activity of C-terminal sub-domain of Prd-DNA-binding domain (Xu et al., 1995) and in the presence of N-terminal sub-domain, C-terminal sub-domain is dispensable for the regulation of downstream genes in Drosophila (Cai et al., 1994). Successive studies with Pax6-DNA complex and Pax5-DNA complex uncover the involvement of C-terminal sub-domain in DNA-binding activity. It was observed that 10 residues from N-terminal α-helical region, 8 residues from N-terminal β-hairpin structure region, 11 residues from linker region, and 9 residues from C-terminal α-helical region of Pax6, directly or via water molecule interact with the DNA and high degree of sequence homology C-subdomain across different Pax family members supports DNA binding functionality of C-subdomain of all Pax family member proteins. (Xu et al., 1999; Garvie et al., 2001).

*In silico* analysis predicted that the mutant protein is quite stable in the cell but mutant protein is rigid and the solvent accessible surface is also reduced which may lead to reduced DNA and protein binding ability of mutant protein. Subsequent analysis also predict that p.Cys112Trp mutation can also disrupt DNA-Protein and Protein-protein interaction.

Out of 33 mutations identified in human PAX9 gene, 4 missense mutations (p.Thr80Ala, p.Ile87Phe, p.Lys97Glu, and p.Tyr143Cys) and one nonsense mutation (p.Lys114X) within the C-subdomain and one frameshift mutation (c.219insG) that disrupt C-subdomain of PAX9 have been known to be associated with congenital tooth agenesis in human (Stockton et al., 2000; Nieminen et al., 2001; Das et al., 2003; Kapadia et al., 2006a; Bergendal et al., 2011). Pax9 protein shows high affinity for 2 putative paired domain recognition sequences namely, e5 and CD19-2(A-ins). Those two probes are often used to study the DNA binding activity by Electrophoretic Mobility Shift Assay (EMSA). For p.Ile87Phe and p.Lys97Glu change EMSA experiments showed reduced affinity for those probes, indicating the involvement of C-terminal subdomain in DNA-binding activity (Kapadia et al., 2006b; Wang et al., 2009b). Beside sequence specific DNA-binding, Pax9 protein was proven to interact with another transcription factor Msx1 protein and together they activate the expression of Bmp4. Protein-protein interaction between Pax9 and Msx1 was used as an experimental model to study the effect of mutation in PAX9 on protein-protein interaction. A number of PAX9 mutations were studied regarding the protein-protein interaction between Pax9 and Msx1; most of the mutants were able to interact with Msx1, although with variable affinity (Wang et al., 2009a).

In this study functional characterizations of the identified p.Cys112Trp variation using *in vitro* DNA-protein and protein-protein interaction were also carried out. DNA-protein interaction through EMSA using e5 probe revealed that this variation disrupt the affinity of mutant protein for putative paired domain binding site. In addition to DNA-binding activity, affinity of mutant protein for Msx1 protein was also very low as it was observed in GST-pull down assay. Taken together our data suggests that due to p.Cys112Trp substitution overall biological activity in terms of DNA-binding and protein-protein interaction was largely compromised.

In this study it was observed that wild-type *PAX9* up-regulates the expression of both *WNT4* and *WN7b* but c.336C>G change, that reduces the DNA binding activity of *PAX9*, unable to up-regulate those two genes in HEK293 cell line. It is worth mentioning here that a few studies on regulation of Wnt signalling by other members of *Pax* gene family support our finding. Torban *et al*. (2006) demonstrate that during kidney development expression of *WNT4* is regulated by *PAX2*. They also experimentally demonstrate the ability of Pax2 protein to interact with DNA probe derived from *WNT4* promoter (Torban et al., 2006). Another observation by Nornes *et al*. (1996) supports the plausible regulation of *WNT4* expression by *PAX9*; in their study they demonstrated that zebrba fish paired domain of both Pax2 and Pax9 have similar binding affinity for 5 different paired domain binding DNA sequences (Nornes et al., 1996). Taking together, it can be concluded that both the *WNT4* and *WNT7b* are the novel targets of *PAX9*. However, it will requires more experimental evidence to decipherer the actual interaction of *PAX9* and those two *WNT* genes during the process of mammalian tooth development in future.

To dissect out the molecular basis of such impairment *in silico* studies were carried out. *In silico* analysis suggests that p.Cys112Trp variation is located within the loop between α-helix 2 and α-helix 3 and may play crucial role in the proper orientation of those two α-helices. Further study using homology modelling revealed that when Cys is substituted with Trp at p.112 position the overall distance between 112Trp and 83Val located within the first α-helix of C-subdomain becomes less compared to 112Cys and 83Val. Tryptophan, which is a bulky amino acid may cause steric hindrance of its neighbouring amino acids that might subsequently alter the structural organization of C-terminal subdomain leading to impairment of DNA-protein and also to some extant protein-protein interaction. It was thus inferred that loss of function of mutant PAX9 due to p.Cys112Trp substitution was associated with impaired tooth development leading to posterior tooth agenesis in DEN14 family.

## 5 Acknowledgments

We are thankful to all the members of the DEN14 family and all control individuals who provided their valuable time to participate in this study. Prof. S. Ganesh, Department of Biological Science and Bioengineering, IIT Kanpur, India, for his kind gift of pCDNA3.1(+) Myc-His(A) vector. We are grateful to Department of Biotechnology, Govt. of India for providing funds for this study. TS acknowledge the receipt of Junior and Senior Research Fellowship from University Grants Commission, Govt. of India (http://www.ugc.ac.in/).

## 6 Conflict of Interest

This study was supported by The Department of Biotechnology, Govt. of India (Grant Number: BT/PR12638/MED/12/467/2009 dated18/03/2010;http://dbtindia.nic.in/index.asp). The authors do not have any financial relationship and declare no conflict of interest.

## 8 Author Contribution

P.D. and T.S. conceived and planned the experiments. S.K. identified and performed clinical investigations. A.G. planed the experiment for GST pull down. P.D. supervised and arranged funding. P.R. did bioinformatics analysis. T.S. performed all the experiments. P.D. and T.S. wrote the manuscript. T.S., S.K., A.G. and P.D. critically review to the findings and the manuscript

## Notes

Funding information This study was supported by The Department of Biotechnology, Govt. of India (Grant Number: BT/PR12638/MED/12/467/2009 dated 18/03/2010; http://dbtindia.nic.in/index.asp).

### Competing Interest Statement

The authors have declared no competing interest.

## References

Adzhubei, I.A., Schmidt, S., Peshkin, L., Ramensky, V.E., Gerasimova, A., Bork, P., et al., 2010. A method and server for predicting damaging missense mutations. Nat Methods 7, 248–9. http://10.1038/nmeth0410-248.

Alfawaz, S., Fong, F., Plagnol, V., Wong, F.S., Fearne, J., Kelsell, D.P., 2013. Recessive oligodontia linked to a homozygous loss-of-function mutation in the SMOC2 gene. Arch Oral Biol 58, 462–6. http://10.1016/j.archoralbio.2012.12.008.

Arte, S., Parmanen, S., Pirinen, S., Alaluusua, S., Nieminen, P., 2013. Candidate gene analysis of tooth agenesis identifies novel mutations in six genes and suggests significant role for WNT and EDA signaling and allele combinations. PLoS One 8, e73705. http://10.1371/journal.pone.0073705.

Bergendal, B., Klar, J., Stecksen-Blicks, C., Norderyd, J., Dahl, N., 2011. Isolated oligodontia associated with mutations in EDARADD, AXIN2, MSX1, and PAX9 genes. Am J Med Genet A 155a, 1616–22. http://10.1002/ajmg.a.34045.

Bonczek, O., Balcar, V.J., Sery, O., 2017. PAX9 gene mutations and tooth agenesis: A review. 92, 467–476. http://10.1111/cge.12986.

Cai, J., Lan, Y., Appel, L.F., Weir, M., 1994. Dissection of the Drosophila paired protein: functional requirements for conserved motifs. Mech Dev 47, 139–50.

Das, P., Hai, M., Elcock, C., Leal, S.M., Brown, D.T., Brook, A.H., et al., 2003. Novel missense mutations and a 288-bp exonic insertion in PAX9 in families with autosomal dominant hypodontia. Am J Med Genet A 118a, 35–42. http://10.1002/ajmg.a.10011.

De Coster, P.J., Marks, L.A., Martens, L.C., Huysseune, A., 2009. Dental agenesis: genetic and clinical perspectives. J Oral Pathol Med 38, 1–17. http://10.1111/j.1600-0714.2008.00699.x.

Dugan, S.L., Temme, R.T., Olson, R.A., Mikhailov, A., Law, R., Mahmood, H., et al., 2015. New recessive truncating mutation in LTBP3 in a family with oligodontia, short stature, and mitral valve prolapse. Am J Med Genet A 167, 1396–9. http://10.1002/ajmg.a.37049.

Garvie, C.W., Hagman, J., Wolberger, C., 2001. Structural studies of Ets-1/Pax5 complex formation on DNA. Mol Cell 8, 1267–76.

G.C.P van Zundert, J.P.G.L.M. Rodrigues, M. Trellet, C. Schmitz, P.L. Kastritis, E. Karaca, A.S.J. Melquiond, M. van Dijk, S.J. de Vries and A.M.J.J. Bonvin (2016). “The HADDOCK2.2 webserver: User-friendly integrative modeling of biomolecular complexes.” J. Mol. Biol., 428, 720–725 (2015).

Hubbard, R. E., & Kamran Haider, M. (2010). Hydrogen Bonds in Proteins: Role and Strength. Encyclopedia of Life Sciences. doi:10.1002/9780470015902.a0003011.pub2.

Hess, et al. (2008) GROMACS 4:L Algorithms for Highly Efficient, Load-Balanced, and Scalable Molecular Simulation J. Chem. Theory Comput. 4: 435–447.

Kantaputra, P., Sripathomsawat, W., 2011. WNT10A and isolated hypodontia. Am J Med Genet A 155a, 1119–22. http://10.1002/ajmg.a.33840.

Kantaputra, P.N., Kaewgahya, M., Hatsadaloi, A., Vogel, P., Kawasaki, K., Ohazama, A., et al., 2015. GREMLIN 2 Mutations and Dental Anomalies. J Dent Res 94, 1646–52. http://10.1177/0022034515608168.

Kapadia, H., Frazier-Bowers, S., Ogawa, T., D’Souza, R.N., 2006a. Molecular characterization of a novel PAX9 missense mutation causing posterior tooth agenesis. Eur J Hum Genet 14, 403–9. http://10.1038/sj.ejhg.5201574.

Kapadia, H., Frazier-Bowers, S., Ogawa, T., D’Souza, R.N., 2006b. Molecular characterization of a novel PAX9 missense mutation causing posterior tooth agenesis. Eur J Hum Genet 14, 403–409.

Lammi, L., Arte, S., Somer, M., Jarvinen, H., Lahermo, P., Thesleff, I., et al., 2004. Mutations in AXIN2 cause familial tooth agenesis and predispose to colorectal cancer. Am J Hum Genet 74, 1043–50. http://10.1086/386293.

Laskowski, R. A., Jabłońska, J., Pravda, L., Vařeková, R. S., & Thornton, J. M. (2018). PDBsum: Structural summaries of PDB entries. Protein science : a publication of the Protein Society, 27(1), 129–134. https://doi.org/10.1002/pro.3289.

Liam J. McGuffin, Kevin Bryson, David T. Jones, The PSIPRED protein structure prediction server, Bioinformatics, Volume 16, Issue 4, April 2000, Pages 404–405, https://doi.org/10.1093/bioinformatics/16.4.404

Lyskov S, Gray JJ (2008) The RosettaDock server for local protein-protein docking. Nucleic Acids Research 36 https://doi.org/10.1093/nar/gkn216

Massink, M.P., Creton, M.A., Spanevello, F., Fennis, W.M., Cune, M.S., Savelberg, S.M., et al., 2015. Loss-of-Function Mutations in the WNT Co-receptor LRP6 Cause Autosomal-Dominant Oligodontia. Am J Hum Genet 97, 621–6. http://10.1016/j.ajhg.2015.08.014.

Mensah, J.K., Ogawa, T., Kapadia, H., Cavender, A.C., D’Souza, R.N., 2004. Functional analysis of a mutation in PAX9 associated with familial tooth agenesis in humans. J Biol Chem 279, 5924–33. http://10.1074/jbc.M305648200.

Molin, A., Lopez-Cazaux, S., Pichon, O., Vincent, M., Isidor, B., Le Caignec, C., 2015. Patients with isolated oligo/hypodontia caused by RUNX2 duplication. Am J Med Genet A 167, 1386–90. http://10.1002/ajmg.a.37052.

Nieminen, P., Arte, S., Tanner, D., Paulin, L., Alaluusua, S., Thesleff, I., et al., 2001. Identification of a nonsense mutation in the PAX9 gene in molar oligodontia. Eur J Hum Genet 9, 743–6. http://10.1038/sj.ejhg.5200715.

Nornes, S., Mikkola, I., Krauss, S., Delghandi, M., Perander, M., Johansen, T., 1996. Zebrafish Pax9 encodes two proteins with distinct C-terminal transactivating domains of different potency negatively regulated by adjacent N-terminal sequences. J Biol Chem 271, 26914–23.

Ogawa, T., Kapadia, H., Feng, J.Q., Raghow, R., Peters, H., D’Souza, R.N., 2006. Functional consequences of interactions between Pax9 and Msx1 genes in normal and abnormal tooth development. J Biol Chem 281, 18363–9. http://10.1074/jbc.M601543200.

Ogawa, T., Kapadia, H., Wang, B., D’Souza, R.N., 2005. Studies on Pax9-Msx1 protein interactions. Arch Oral Biol 50, 141–5. http://10.1016/j.archoralbio.2004.09.011.

Polder, B.J., Van’t Hof, M.A., Van der Linden, F.P., Kuijpers-Jagtman, A.M., 2004. A meta-analysis of the prevalence of dental agenesis of permanent teeth. Community Dent Oral Epidemiol 32, 217–26. http://10.1111/j.1600-0528.2004.00158.x.

Ranjan P N, Devi C, Das P. Bioinformatics analysis of SARS-CoV-2 RBD mutant variants and insights into antibody and ACE2 receptor binding. bioRxiv; 2021. DOI: 10.1101/2021.04.03.438113.

Prashant Ranjan, Bhagyalaxmi Mohapatra, Parimal Das et al. A rational drug designing: What bioinformatics approach tells about the wisdom of practicing traditional medicines for screening the potential of Ayurvedic and natural compounds for their inhibitory effect against COVID-19 Spike, Indian strain Spike, Papain-like protease and Main Protease protein, 20 May 2020, PREPRINT (Version 1) available at Research Square [https://doi.org/10.21203/rs.3.rs-30366/v1]

Sambrook, J., Russell, D.W., 2001. Molecular Cloning: A Laboratory Manual, Cold Spring Harbor Laboratory Press.

Sarkar, T., Bansal, R., Das, P., 2014. Whole genome sequencing reveals novel non-synonymous mutation in ectodysplasin A (EDA) associated with non-syndromic X-linked dominant congenital tooth agenesis. PLoS One 9, e106811. http://10.1371/journal.pone.0106811.

Schwarz, J.M., Cooper, D.N., Schuelke, M., Seelow, D., 2014. MutationTaster2: mutation prediction for the deep-sequencing age. Nat Methods 11, 361–2. http://10.1038/nmeth.2890.

Stockton, D.W., Das, P., Goldenberg, M., D’Souza, R.N., Patel, P.I., 2000. Mutation of PAX9 is associated with oligodontia. Nat Genet 24, 18–9. http://10.1038/71634.

Shula Shazman, Gershon Celniker, Omer Haber, Fabian Glaser, Yael Mandel-Gutfreund (2007) Patch Finder Plus (PFplus): A web server for extracting and displaying positive electrostatic patches on protein surfaces. Nucleic Acids Res., 35:526–30.

Tao, R., Jin, B., Guo, S.Z., Qing, W., Feng, G.Y., Brooks, D.G., et al., 2006. A novel missense mutation of the EDA gene in a Mongolian family with congenital hypodontia. J Hum Genet 51, 498–502. http://10.1007/s10038-006-0389-2.

Torban, E., Dziarmaga, A., Iglesias, D., Chu, L.L., Vassilieva, T., Little, M., et al., 2006. PAX2 activates WNT4 expression during mammalian kidney development. J Biol Chem 281, 12705–12. http://10.1074/jbc.M513181200.

van den Boogaard, M.J., Creton, M., Bronkhorst, Y., van der Hout, A., Hennekam, E., Lindhout, D., et al., 2012. Mutations in WNT10A are present in more than half of isolated hypodontia cases. J Med Genet 49, 327–31. http://10.1136/jmedgenet-2012-100750.

Vastardis, H., Karimbux, N., Guthua, S.W., Seidman, J.G., Seidman, C.E., 1996. A human MSX1 homeodomain missense mutation causes selective tooth agenesis. Nat Genet 13, 417–21. http://10.1038/ng0896-417.

Wang, Y., Groppe, J.C., Wu, J., Ogawa, T., Mues, G., D’Souza, R.N., et al., 2009a. Pathogenic mechanisms of tooth agenesis linked to paired domain mutations in human PAX9. Hum Mol Genet 18, 2863–74. http://10.1093/hmg/ddp221.

Wang, Y., Wu, H., Wu, J., Zhao, H., Zhang, X., Mues, G., et al., 2009b. Identification and functional analysis of two novel PAX9 mutations. Cells Tissues Organs 189, 80–7. http://10.1159/000151448.

Xu, H.E., Rould, M.A., Xu, W., Epstein, J.A., Maas, R.L., Pabo, C.O., 1999. Crystal structure of the human Pax6 paired domain-DNA complex reveals specific roles for the linker region and carboxy-terminal subdomain in DNA binding. Genes Dev 13, 1263–75.

Xue LC, Rodrigues JP, Kastritis PL, Bonvin AM, Vangone A. PRODIGY: a web server for predicting the binding affinity of protein-protein complexes. Bioinformatics. 2016 Dec 1;32(23):3676–3678. doi: 10.1093/bioinformatics/btw514. Epub 2016 Aug 8. PMID: 27503228.

Xu, W., Rould, M.A., Jun, S., Desplan, C., Pabo, C.O., 1995. Crystal structure of a paired domain-DNA complex at 2.5 A resolution reveals structural basis for Pax developmental mutations. Cell 80, 639–50.

Yu, P., Yang, W., Han, D., Wang, X., Guo, S., Li, J., et al., 2016. Mutations in WNT10B Are Identified in Individuals with Oligodontia. Am J Hum Genet 99, 195–201. http://10.1016/j.ajhg.2016.05.012.

